# Mechanobiochemical finite element model to analyze impact-loading-induced cell damage, subsequent proteoglycan loss, and anti-oxidative treatment effects in articular cartilage

**DOI:** 10.1101/2024.07.03.601852

**Authors:** Joonas P. Kosonen, Atte S.A. Eskelinen, Gustavo A. Orozco, Mitchell C. Coleman, Jessica E. Goetz, Donald D. Anderson, Alan J. Grodzinsky, Petri Tanska, Rami K. Korhonen

## Abstract

Joint trauma often leads to articular cartilage degeneration and post-traumatic osteoarthritis (PTOA). Pivotal determinants include trauma-induced excessive tissue strains that damage cartilage cells. As a downstream effect, these damaged cells can trigger cartilage degeneration via oxidative stress, cell death, and proteolytic tissue degeneration. N-acetylcysteine (NAC) has emerged as antioxidant capable of inhibiting oxidative stress, cell death, and cartilage degeneration post-impact. However, temporal effects of NAC are not fully understood and remain difficult to assess solely by physical experiments. Thus, we developed a computational framework to simulate a drop tower impact of cartilage with finite element analysis in ABAQUS, and model subsequent oxidative stress-related cell damage, and NAC treatment upon cartilage proteoglycan content in COMSOL Multiphysics, based on prior *ex vivo* experiments. Model results provide evidence that by inhibiting further cell damage by mechanically induced oxidative stress, immediate NAC treatment can reduce proteoglycan loss by mitigating cell death (loss of proteoglycan biosynthesis) and enzymatic proteoglycan depletion. Our simulations also indicated that delayed NAC treatment may not inhibit cartilage proteoglycan loss despite reduced cell death after impact. These results enhance understanding of temporal effects of impact-related cell damage and treatment that are critical for the development of effective treatments for PTOA. In the future, our modeling framework could increase understanding of time-dependent mechanisms of oxidative stress and downstream effects in injured cartilage and aid in developing better treatments to mitigate PTOA progression.

**Author summary:** Post-traumatic osteoarthritis is a debilitating disease which is often initiated by trauma and characterized by cartilage degeneration. The degeneration is partly driven by trauma-induced damage, including oxidative stress, to cartilage cells. Multiple drugs have been studied to counteract the cell damage, but it remains difficult to inhibit the disease progression since temporal effects of such treatments are not fully understood. Here, we developed a computational framework to study effects of antioxidant treatment in mechanically impacted cartilage and compared our computational simulations with previously published experiments. Our results showed that the high strain induced cell damage occurring in the mechanically impacted region could be mitigated by N-acetylcysteine treatment. This mechanism could partly explain reduced cartilage proteoglycan loss compared to untreated samples. Our modeling framework could help enhance development of treatments to better inhibit osteoarthritis progression.

## 1. Introduction

Disturbances in articular joint homeostasis after traumatic injuries, such as intra-articular fractures and anterior cruciate ligament rupture, can ultimately lead to post-traumatic osteoarthritis (PTOA) characterized by pain, stiffness, and cartilage degeneration [1–3]. Although the mechanisms of PTOA onset are not fully understood, several studies have shown that chondrocytes (cartilage cells) play a key role in cartilage health after impact [1,4–6]. The cell-driven degeneration of the tissue has been suggested to be triggered by inflammation [6] and trauma-related alterations in cell mechanotransduction due to excessive tissue strains or strain rates [4,5,7]. The excessive mechanical strains can deform and damage the cells in cartilage, with a downstream effect of promoting excessive production of reactive oxygen species (ROS) that cause oxidative stress [5,8]. Damaged cells experiencing oxidative stress can undergo cell death, whether through necrosis or apoptosis, which decreases biosynthesis of proteoglycan molecules in the extracellular matrix (ECM) [9,10]. In addition, excessive amounts of ROS in damaged cells can act as secondary messengers that upregulate the release of proteolytic enzymes, such as aggrecanase and collagenase enzymes, that further increase the loss of the ECM components [11,12].

Orally or intra-articularly delivered antioxidants can reduce the amount of ROS directly by scavenging ROS and indirectly by fortifying the natural antioxidative system within chondrocytes [11,13,14]. Several antioxidants have been studied [13,15], and N-acetylcysteine (NAC) has emerged as one of the most potent antioxidants in inhibiting cell death and reducing ECM degeneration after *ex vivo* impact [16–20]. NAC has also been shown to inhibit cell death and ECM degeneration in a large animal model after severe impact-injury when administered intra-articularly post-impact [20]. However, the rapid clearance of small NAC molecules (163 Da) from intra-articular space [21] makes it challenging to find an optimized dose sufficient to inhibit acute cell damage and later cartilage degeneration *in vivo* after impact.

Computational models have proved useful for estimating cartilage response to injurious loading, progression of cartilage degeneration, and treatment effects. Biomechanical finite element models with stress/strain-based degeneration algorithms have been used for simulating tissue degeneration under physiologically relevant loading conditions [22–25]. Recently, these biomechanical degeneration models have been coupled with biological mechanisms, such as time-dependent diffusion of pro-inflammatory cytokines [26,27], biomechanically triggered ROS overproduction [28], cell death [29], and enzymatic degeneration of proteoglycans [26]. Building on these mechanobiological degeneration models, recent work has simulated the effects of anti-inflammatory treatments on time-dependent cartilage degeneration [30,31]. Yet, there are no models combining injurious loading, oxidative stress-induced cell damage, and damaged cell-driven degeneration of cartilage with computational simulation of antioxidant treatment aiming to prevent cellular oxidative injury following a high-energy impact. Such models could be used to explore why an intra-articular anti-oxidative injection may or may not work in any given clinically relevant scenario (type of injury, treatment timing, dosage, etc.).

In this study, we developed a new computational modeling framework to simulate a high energy impact, impact-induced oxidative cell damage, acute cell death and adaptation of proteoglycan content. This framework was then used to simulate the effects of short-term NAC treatment in mature bovine cartilage, with an aim to understand which mechanobiologically relevant modeling parameters can explain the cell viability and proteoglycan content quantitatively determined in prior experiments [17]. We hypothesized that i) higher proteoglycan content in immediately NAC treated vs. untreated samples after impact could be partly explained by inhibited strain-induced local cell oxidative damage, proteolytic degeneration of proteoglycans, and cell death, and that ii) although impact-induced oxidation driven cell death could be inhibited with NAC treatment delivered at a later time point after mechanical impact, delayed treatment is insufficient to fully protect matrix proteoglycan content due to substantial release of proteolytic enzymes before the treatment. Due to a lack of accurate and repeatable measures of cell-level parameters such as cell death rate and chondrocyte protection rate by NAC, we investigated the effect of the most important parameters on cell death and proteoglycan content via sensitivity analysis. This approach marks an important step towards understanding and numerically estimating the underlying mechanobiological effects of antioxidant treatment at cell and tissue levels in cartilage after severe impact-injury. Ultimately, our modeling framework could help in designing more efficient treatments to mitigate PTOA.

## 2. Methods

### 2.1. Previous experiments as a basis of the computational framework

Our computational modeling framework leverages prior *ex vivo* drop-tower impact experiments of mature bovine osteochondral plugs (25 mm wide) conducted by Martin et al. [17] (Fig 1A). The following experimental data were compared against outputs from our computational model: **1)** cell viability in untreated samples 200 µm from the impacted surface (confocal microscopy) 1-, 3-, 6-, 12-, 24-, and 48-hours after impact (Fig. 1A and B), **2)** cell viability in samples treated with NAC (2mM) at day 2 with 0-, 1-, 4-, and 12-hour treatment delay (Fig. 1C), and **3)** relative proteoglycan content at days 7 and 14 post-impact in samples with and without immediate 1-day NAC treatment (Fig. 1D). In the experiments, the relative proteoglycan content was quantified with dimethyl methylene blue assay of 4-mm wide impacted vs. intact regions dissected from original samples [17].

**Fig. 1.**
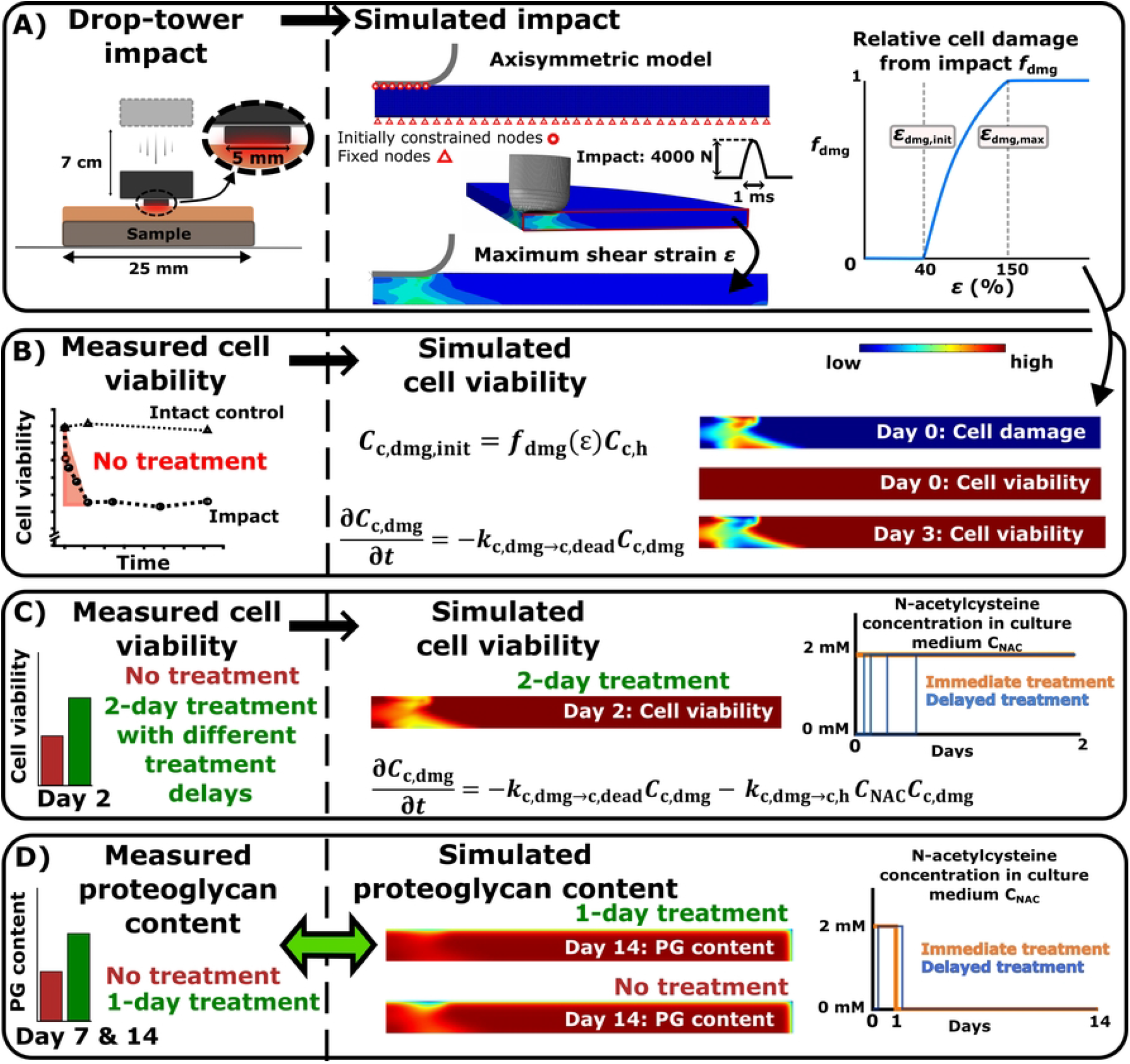
Workflow. A) The biomechanical response of cartilage to drop-tower impact was first simulated based on prior experiments. Maximum shear strains were computed at the time of peak impact force (4000N) to identify cell damage (i.e., cells experiencing oxidative stress). B) Time-dependent loss of cell viability from impact was next simulated for comparison with experimental findings. C) Then N-acetylcysteine (NAC)-induced cell recovery after 0, 1, 4, and 12-hour post-impact treatment delay was simulated, and viability was matched with experiments. D) Finally, proteoglycan degeneration driven by the damaged cells and the mitigating effect of NAC treatment was simulated.

### 2.2. Finite element model for simulating impact loading and shear strain-triggered cell damage

An axisymmetric finite element model of the cartilage (width of 12.5 mm, height of 1 mm) and a beveled flat-ended indenter (5 mm diameter with 1 mm circle radius for the rounded edge) was constructed to simulate the impact. Cartilage was modeled as a fibril-reinforced poroviscoelastic material with Donnan osmotic swelling [32,33] while the indenter was assumed rigid. The depth-dependent content and structure (i.e. proteoglycan content, water content, collagen density and orientation, and material parameters of the model) were estimated based on prior reports of mature bovine cartilage (see Supplementary material S1) [32–34]. As an initial condition in the model, cartilage was allowed to swell until mechanical equilibrium (physiological salt concentration, i.e., 0.15 M NaCl) [32] followed by simulation of the cartilage-indenter impact where sinusoidal-like force was applied on the indenter. Earlier drop-tower studies have reported sinusoidal-like force-response to impact over 0.6-2 ms [35–37], thus we assumed impact time *t*_impact_= 1 ms (Fig. 1A). With this impact time we calculated the average impact force *F*_impact,av_ = *mv*/*t*_impact_, where impact velocity 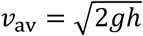, m is mass of the impactor, *g* is the gravity constant, *h* is drop height of the impactor) to be *F*_impact,av_ = 2350 N (average stress *σ*_impact,*av*_ = 120 MPa) and corresponding peak impact force to be *F*_impact,peak_ = 4000 N (Fig 1A; peak impact stress *σ*_peak_ = 200 MPa). We consider these peak impact force estimates reasonable, since they are in similar range as earlier drop-tower impact experiments showing 50-70 MPa peak impact stresses (800-1100 N peak impact forces) with an indenter of 5.5 mm in diameter [38]. However, sensitivity analysis with 2000N (100MPa peak impact stress) and 6000N peak impact forces (300MPa peak impact stress) was also conducted to analyze effect of the impact force on maximum shear strains and cell damage (See supplementary material S2).

Since high local strains trigger oxidative stress and cell death [5], we calculated the maximum shear strain from the Green-Lagrangian strain tensor at the peak impact force (0.5 ms). The maximum shear strain distribution was used to define the initial cell damage (day 0 initial condition) with a non-linear cellular damage function *f*_dmg_(*ε*) [23]:

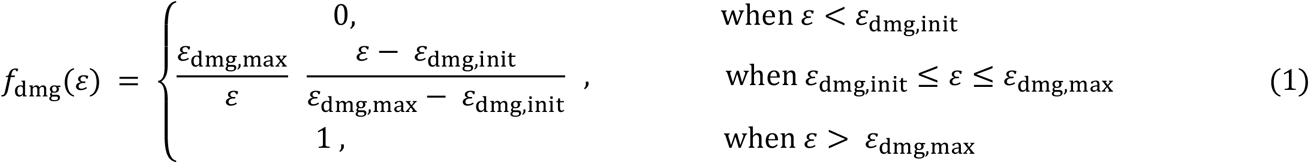

where *ε* is the maximum shear strain, *ε*_dmg,init_ = 40% is the strain threshold describing the initiation of cell damage, and *ε*_dmg,max_ = 150% is the maximum cell damage threshold describing the limit when all cells are damaged [7]. This initial cell damage was used as an input for the cell viability and proteoglycan content simulations (see section 2.3, Eq. 6).

The biomechanical model was constructed in ABAQUS (v. 2023, Dassault Systèmes, Providence, RI, USA) with 2506 continuum pore pressure elements (type: CAX4P) and solved with transient soils analysis. Indenter-cartilage contact was modeled using tabular pressure-overclosure relationship. Nodes of the cartilage geometry initially in contact with the indenter were constrained in the radial direction during the simulations. The bottom surface of the cartilage tissue was fixed in axial and radial directions (osteochondral plug in Martin et al. [17]; bone was not included in the model). Fluid flow and radial movement of the nodes was prevented on the symmetry axis of the plug (axisymmetric boundary condition). Fluid flow was allowed through the free boundaries. Mesh convergence was verified (see Supplementary material S3).

### 2.3. Modeling cell viability, proteoglycan degeneration, and NAC treatment

In our modeling framework, we simulated distributions of healthy, damaged and dead cells, where initially healthy cells were turned into damaged cells due to excessive local strains caused by the impact (Eq. 1). The damaged cell population was assumed to have less efficient antioxidative defenses, thus they were allowed to die over time due to being more susceptible to oxidative stress after the impact [39–41]. In the model, the cells in a damaged state released proteolytic enzymes that could degenerate proteoglycans and no purely mechanically induced degeneration was considered. The total biosynthesis of proteoglycans in cartilage was decreased due to lower number of viable cells (healthy + damaged). The cell damage was mitigated by diffusion of NAC from the free tissue boundaries, inhibiting the downstream effects of cell damage and restoring the antioxidative defense system of the cells rendering them back into healthy cellular state [20].

All cell-related processes were modeled in Comsol Multiphysics (v. 5.6, Burlington, MA, USA) with reaction–diffusion partial differential equations [26,28]:

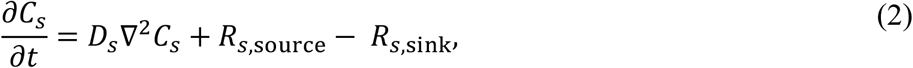

where *t* is time, *C*_*s*_ is the concentration of a given constituent species within cartilage, *D*_*s*_ is the effective diffusion constant, R_*s*,source_ is the source term, and R_*s*,sink_ the sink term of species *s* (*s* = healthy, damaged or dead cell population or proteoglycan, proteolytic enzyme, or NAC concentration) [26]. Effective diffusion of proteolytic enzymes was modeled depth-dependent according to the defined proteoglycan content as in [26]. Source/sink terms for proteoglycans and proteolytic enzymes (describing degeneration of proteoglycans) were modeled according to Michaelis-Menten kinetics as in [26] and [28]. For NAC, we assumed isotropic diffusion as *D*_NAC_ = 120. 10^−6^ m^2^/s based on its molecular weight (163.2 Da) [42]. Due to lack of experimental data regarding NAC half-life and chemical reaction rates in cartilage, source/sink terms for NAC were assumed zero (i.e., NAC was only diffusing into cartilage without further changes until free diffusion of NAC out of cartilage when culture media was changed).

Damaged chondrocytes were allowed to die due to oxidative stress. Unlike in previous studies [28,29], we did not explicitly model ROS concentration in damaged cells due to high variation in reaction kinetics of different ROS molecules [14]. Instead, we implicitly modeled cellular oxidative stress through the concentration of damaged chondrocytes *C*_c,dmg_, that could be further altered by the presence of NAC:

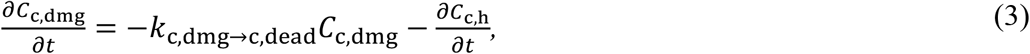

where *k*_c,dmg→c,dead_ is the cell death rate for damaged cells, and *C*_c,h_ is the concentration of healthy cells. Accordingly, the recovery of damaged chondrocytes back to healthy was modeled as

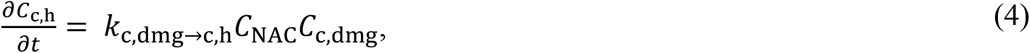

where *k*_c,dmg→c,h_ is the chondrocyte protection rate due to NAC (recovery of damaged cells back to healthy) and *C*_NAC_ is the NAC concentration. Since Martin et al. [17] reported on average less than 5% change in cell viability in unimpacted control samples after 3 days of culture, we did not assume spontaneous, basal chondrocyte death or recovery.

Proteoglycan loss was modeled by simulating increased proteolytic activity observed earlier after impact injury [19,43]. The production of proteolytic enzymes was increased according to an exponential stimulus function *S* (for more details, see Supplementary material S4), which was elevated in the areas of damaged cells *C*_c,dmg_:

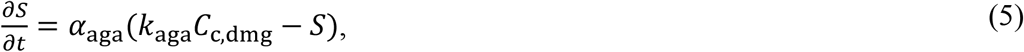

where *α*_aga_ = 0.4. 10−5 s^−1^ is the rate constant for stimulus and *k*_aga_ is a stimulus constant for release from damaged cells. Initial cell stimulus and proteolytic enzyme concentration was set to zero, and zero flux for proteolytic enzymes was set at all boundaries. Initial proteoglycan concentration was calculated from the fixed charge density distribution used in the biomechanical impact model (for more details of the biochemical model, see supplementary material S4) [32,44].

Due to a lack of data on cell distribution of the impacted samples in [17], initial healthy cell concentration was assumed homogenous (*C*_c,h,init_ = 0.5 × 10^14^ m^-3^) [45]. Also, no cell proliferation was considered (*i*.*e*. sum of damaged, healthy, and dead cells was assumed constant *C*_c,dmg_ + *C*_c,h_ + *C*_c,dead_ = *C*_c,h,init_). Assuming a fraction of cells would become damaged after impact (see Eq. 1) [5], we set the initial cell damage as:

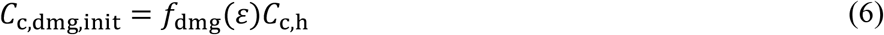

The initial NAC concentration within the cartilage was set to zero, and the concentration on the free surfaces (top and outer surface) was set to 2 mM (Fig. 1C). To simulate the delayed administration of NAC (0, 1, 2, 3, 4 and 12 hours after impact, Fig. 1C), the NAC concentration on the free surfaces was increased from 0 mM to 2 mM with a step function. Radial and axial fluxes through the boundaries for proteoglycan and proteolytic enzyme molecules were defined as described previously by Kar et al. [26].

### 2.5 Sensitivity analysis and reference parameters

Sensitivity analyses were conducted to analyze the effect of relevant model parameters on the cell viability and proteoglycan content (Table 1). These parameters included the maximum cell damage threshold *ε*_dmg,max_, cell death rate for damaged cells *k*_c,dmg→c,dead_, proteolytic enzyme stimulus constant in damaged cells *k*_aga_, and chondrocyte protection rate after NAC treatment *k*_c,dmg→c,h_. Reference parameters were selected so that predicted average cell viability matched experimental outputs [17] (see Section 2.1). Ranges for sensitivity analysis were selected so that predicted average cell viability was within one standard deviation of the experimentally measured mean cell viability. Additionally, we conducted sensitivity analyses for the peak impact force and proteolytic enzyme stimulus constant in damaged cells to study their effect on the initial cell damage and proteoglycan loss (see supplementary material S2 and S5).

**Table 1.** Parameters for sensitivity analysis. Cell viability, proteoglycan degeneration and treatment-related parameters chosen for the sensitivity analysis. Bolded values were chosen as the reference value.

| Parameters | Reference parameters | Range | Description | Reference |
| --- | --- | --- | --- | --- |
| $\varepsilon_{\text{dmg,max}} [\%]$ | **150%** | 100%, **150%**, 200% | Maximum cell damage threshold | [7,17] |
| $k_{\text{c,dmg} \rightarrow \text{c,dead}} [10^{-5} \text{ s}^{-1}]$ | **0.69** | 4.6, **6.9**, 13.9 | Cell death rate for damaged cells | [17] |
| $k_{\text{c,dmg} \rightarrow \text{c,h}} [10^{-4} \text{ m}^3 \text{ mol}^{-1} \text{ s}^{-1}]$ | **0.53** | 0.29, 0.39, **0.53** | Chondrocyte protection rate | [17] |

## 3. Results

### 3.1 By inhibiting cell damage in impacted cartilage, NAC partly reduced proteoglycan loss

With the reference parameters (table 1 in Section 2.5), the mechanical impact model predicted cell damage extending through the full thickness of the tissue, whereas no cell damage was predicted in the non-impacted region (Fig 2A). Without treatment, cell death was observed in the excessively loaded regions of cartilage, and NAC treatment immediately after impact inhibited the acute cell death.

**Fig. 2.**
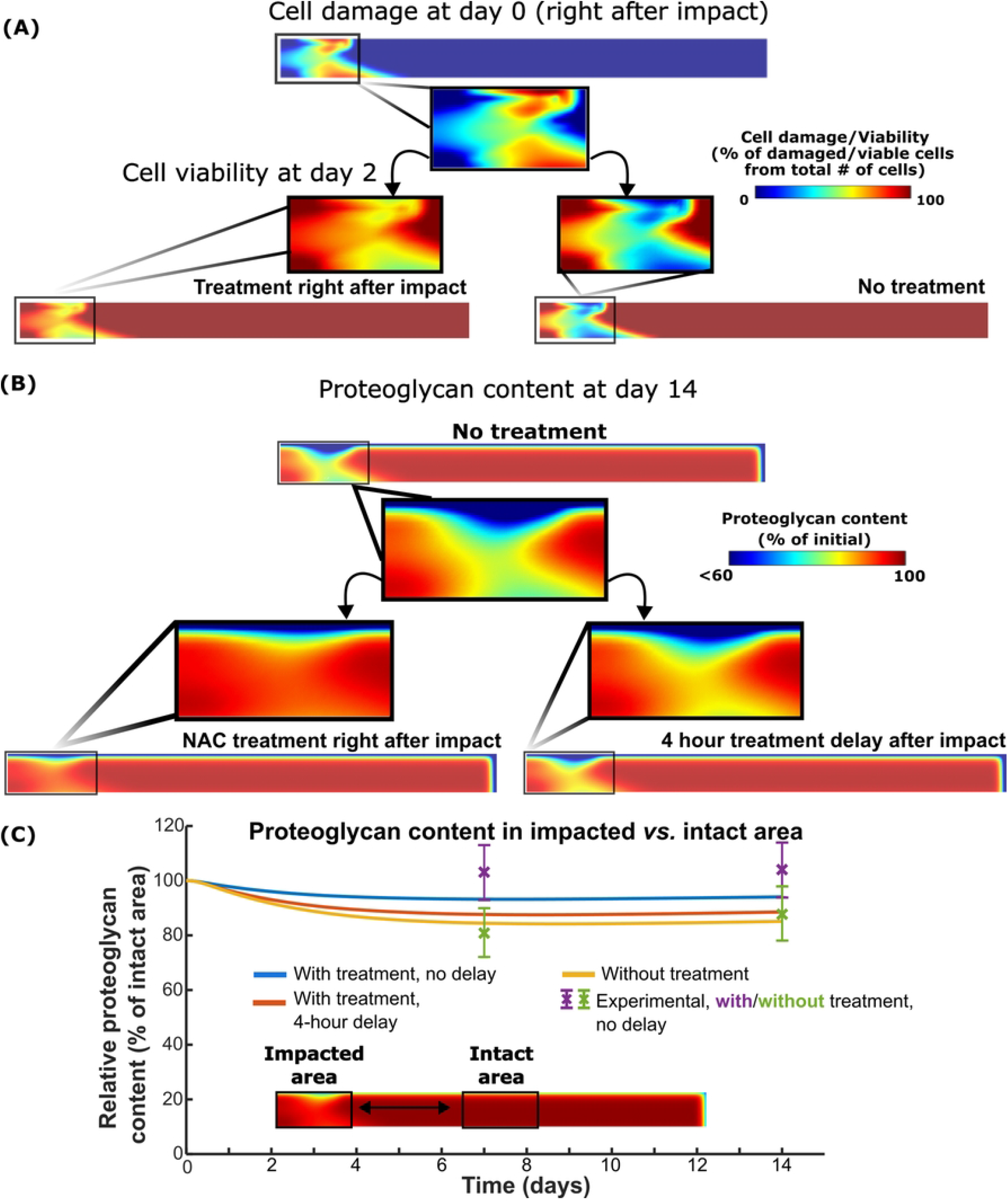
NAC can reduce proteoglycan loss by inhibiting impact-induced cell damage. A) Our simulations showed high cell damage after impact, leading to acute cell death in damaged regions. Immediate post-impact NAC treatment effectively preserved cell viability throughout cartilage depth in the damaged areas. (B)Without NAC treatment, proteoglycan content was subsequently reduced throughout the depth of the cartilage. Immediate post-impact NAC treatment inhibited proteoglycan loss by reducing cell damage, but treatment after a 4-hour delay was less successful in maintaining proteoglycan content. C) Quantitatively, our simulated results matched experiments and showed that 4-hour treatment delay resulted in proteoglycan content resembling the content in untreated cartilage after impact.

The model predicted proteoglycan loss throughout the cartilage depth in the impacted area (Fig. 2B), and the lowest proteoglycan content was located at the superficial zone. Simulated NAC treatment was able to inhibit the proteoglycan content loss in the impacted area when the treatment was administered immediately after the impact loading. Four hours post-impact treatment delay resulted in proteoglycan loss in the superficial and deeper parts of the cartilage closer to that in the untreated samples. Most of the proteoglycan loss was observed within 7 days after impact, and at day 14, the predicted proteoglycan content was 5%, 11% and 14 % lower in the impacted region compared to intact region in immediately treated, 4-hour treatment delay, and without treatment models (Fig. 2C).

### 3.2. Sensitivity analysis of cell damage and predicted cell viability after delayed treatment

Increasing the *ε*_dmg,max_ threshold (Table 1) resulted in lower cell damage in the impacted area (Fig. 3 A-C), while the opposite was observed when decreasing the threshold. That is, *ε*_dmg,max_ = 100% resulted in 65% cell damage for the initially healthy cells in the impacted superficial zone, whereas the reference value *ε*_dmg,max_ = 150% led to 56% and *ε*_dmg,max_ = 200% to 52% cell damage. By selecting *ε*_dmg,max_ = 150% (Fig. 3D), the reference model (with *k*_c,dmg→c,dead_ = 6.9. 10−5 s^-1^, Table 1) replicated well the average cell viability (46%) observed in experiments three days after the impact loading [17]. The reference model also showed the best fit to the experimentally observed cell viability over the entire 3 days after impact, showing 64% cell viability 4 hours after impact (Fig. 3E). In contrast, with *k*_c,dmg→c,dead_ = 13.9. 10−5 s^-1^ and *k*_c,dmg→c,dead_ = 4.6. 10−5 s^-1^, the model overestimated (viability 51% at 4h) and underestimated (viability 72% at 4h) the average cell death, respectively, compared to experiments during the first hours post-impact.

**Fig. 3.**
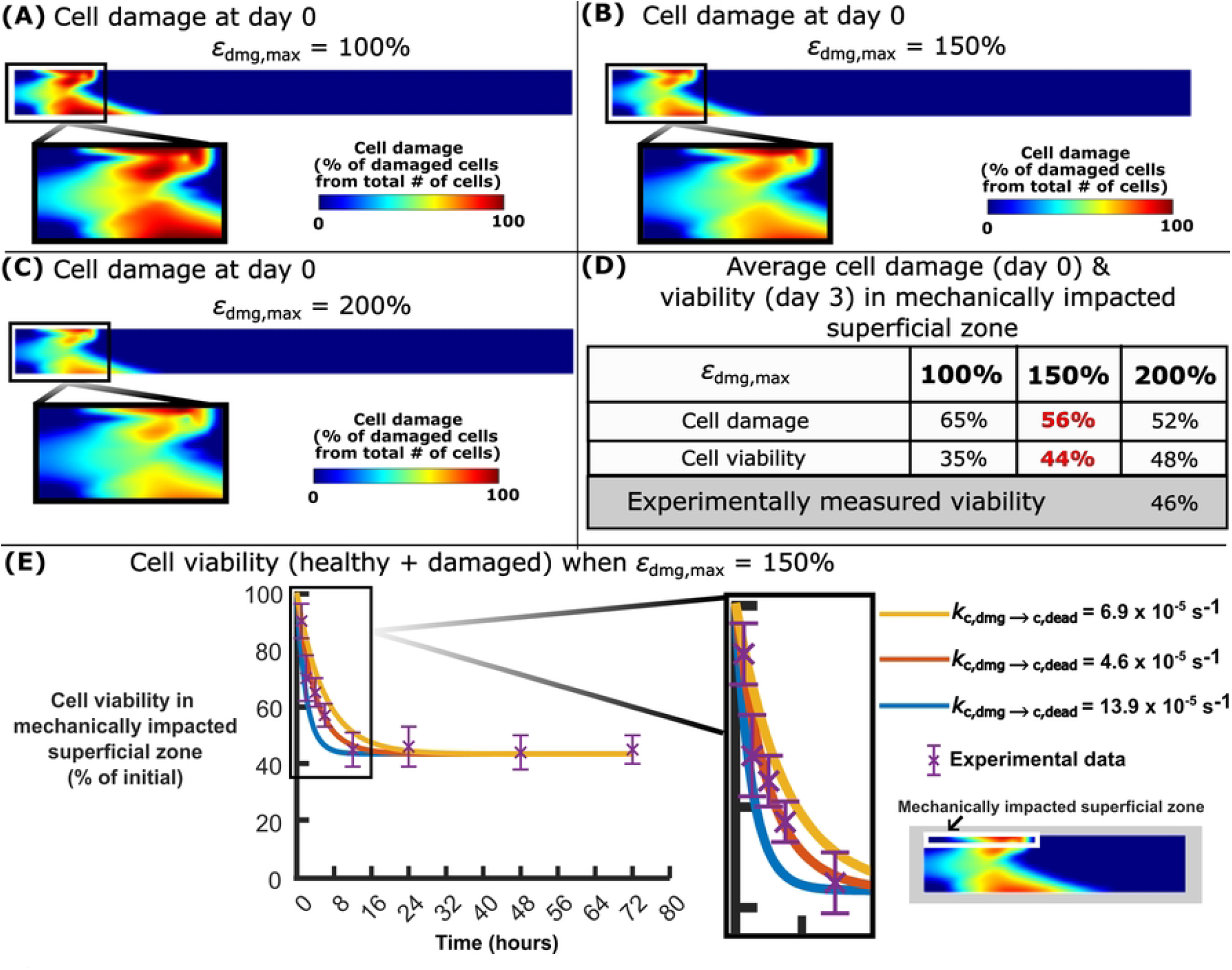
Calibration of cell damage and viability after impact. Cell damage distributions at day 0 after impact, when A) the maximum cell damage threshold *ε*_dmg,max_ = 100 %, B) when *ε*_dmg,max_ = 150 %, and C) when *ε*_dmg,max_ = 200 %. Since D) *ε*_dmg,max_ = 150 % best matched the experiments, E) it was utilized to determine the cell death rate for damaged cells *k*_c,d→c,dead_ over 72 hours after impact. With *k*_c,d→c,dead_ = 6.9 × 10^−5^ s^-1^, the model replicated also the decrease in cell viability (amount of healthy cells maintained) during the first 16 hours after impact. Experimental data show the mean ± standard deviation.

In the case of immediate NAC treatment post-impact, increasing the chondrocyte protection rate (*k*_c,dmg→c,h_) resulted in further improvements in predicted cell viability two days post-impact, especially at the superficial cartilage (Fig. 4A-C). With 2 day NAC treatment and *k*_c,dmg→c,h_ = 0.29. 10^−4^ m^3^mol^-1^s^-1^ in the model, the predicted cell viability was increased from 46% (no NAC treatment) to 69% in the impacted superficial zone (Fig. 4D), while with *k*_c,dmg→c,h_ = 0.39. 10^−4^ m^3^mol^-1^s^-1^ or *k*_c,dmg→c,h_ =0.53. 10^−4^ m^3^mol^-1^s^-1^, the predicted cell viabilities due to treatment were increased to 73% and 77%,respectively. When *k*_c,dmg→h_ = 0.29. 10^−4^ m^3^mol^-1^s^-1^ and *k*_c,dmg→h_ = 0.39. 10^−4^ m^3^mol^-1^s^-1^ was used, the model underestimated cell viability when the treatment was delayed compared to the experiments (data not shown). When *k*_c,dmg→h_ = 0.53. 10^−4^ m^3^mol^-1^s^-1^ was used in the model, cell viabilities were reduced from 77% to 45% as a function of the NAC treatment delay from 0 to 12 hours, respectively (Fig. 4E).

**Fig. 4.**
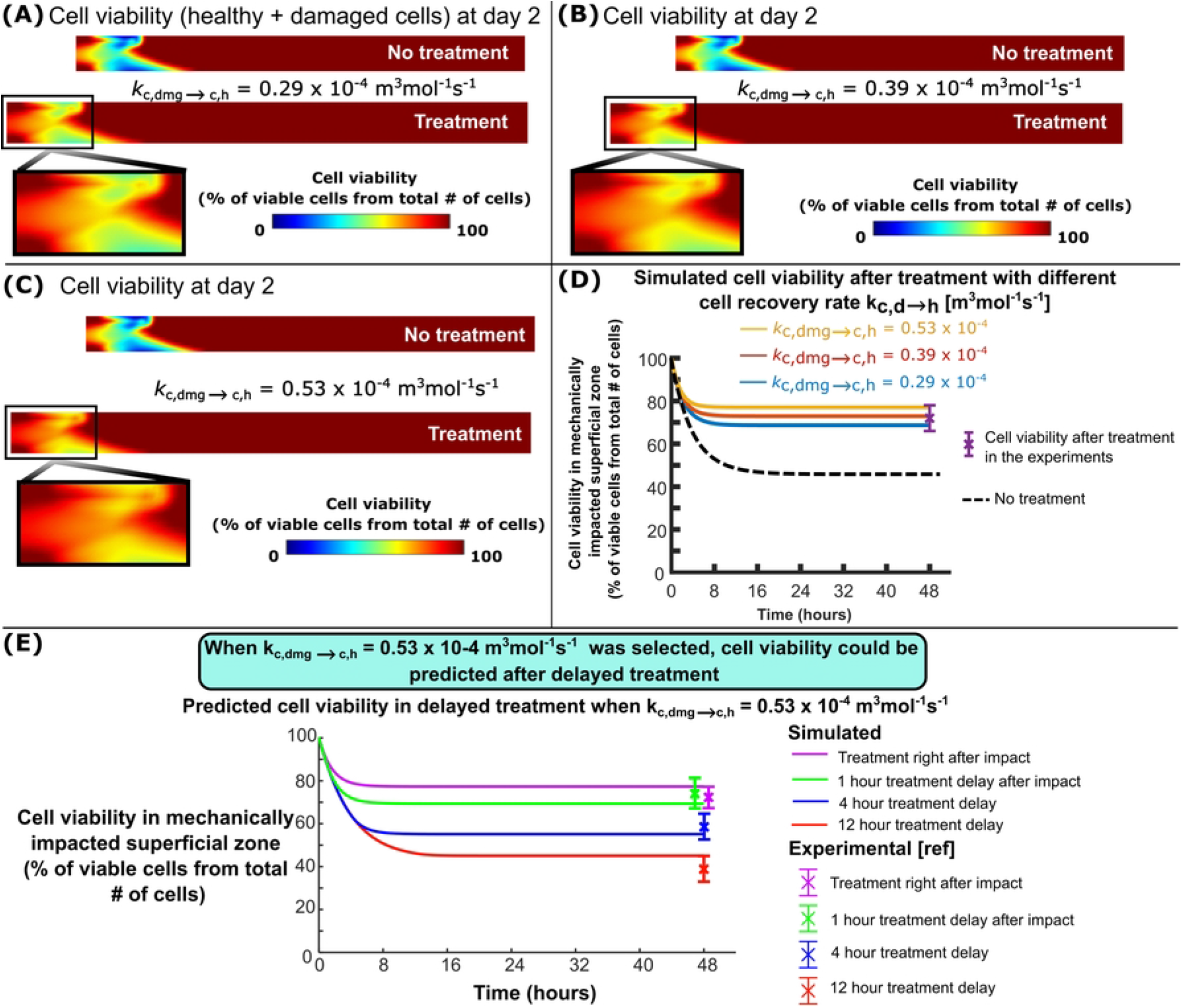
Cell viability after treatment. Cell viability distributions 2 days after impact when A) chondrocyte protection rate *k*_c,dmg→c,h_ = 2.9. 10−5 m^3^mol^-1^s^-1^, B) when *k*_c,dmg→c,h_ = 3.9. 10−5 m^3^mol^-1^s^-1^ and when (B) *k*_c,dmg→c,h_ = 5.3. 10−5 m^3^mol^-1^s^-1^. D) Simulated and experimentally-measured cell viability in the superficial zone of the impacted area over 2 days. E) Comparison of the simulated and experimentally-measured cell viability after 0-, 1-, 4-, and 12-hour treatment delay. With reference parameter *k*_c,dmg→c,h_ = 5.3 × 10^−5^ m^3^mol^-1^s^-1^, our model was able to replicate the cell viability after treatment delay as observed in experiments.

## 4. Discussion

We developed a computational modeling framework to simulate impact induced early-phase cell damage (oxidative stress), cell death, proteoglycan degeneration, and short-term anti-oxidative NAC treatment aiming to counteract the biological effects of the impact. We calibrated the model against quantitative data of cell viability and proteoglycan loss from previous *ex vivo* experiments [17]. The main findings were: i) high shear strains in the impacted region of cartilage can lead to cellular damage and oxidative stress, triggering cell death pathway and proteoglycan degeneration, ii) inhibition of impact-induced oxidative cell damage and cell death with NAC resulted in reduced proteolytic enzyme activity and production (Eq. 5), thereby mitigating proteoglycan loss and iii) since ongoing proteolytic enzyme production was not inhibited, delayed (> 4 hours) treatment may not protect the proteoglycan content despite the fact that cell death could be reduced by NAC.

### 4.1. Simulated cell damage and cell viability after impact

Cell damage was observed in the superficial and deep cartilage due to high shear strains [4,8,46] leading to accumulated cell death over 2 days after impact (Fig 2). This is also consistent with earlier cartilage impact loading experiments that have reported post-impact cell death in superficial and deep cartilage layers [46–49]. Furthermore, our simulations predicted 56% cell damage immediately after impact, in line with the earlier *ex vivo* reports showing harmful ROS production in 60% of superficial chondrocytes 1 hour after impact [47] and 55 % viability (Fig 3D) after 3 days [17]. Similarly, other *ex vivo* impact models have shown that there is a connection between lethal oxidative stress, oxidative stress associated cell damage, and subsequent cell death within an hour after impact [46]. Acute cell death within hours after impact can include different types of cell death such as necrosis and subacute apoptosis [50], both promoted by excessive amounts of ROS [51]. In the current model, net cell death rate considering both necrosis and apoptosis was used to replicate rapid cell death [17] and different cell death types were not considered. However, a large number of necrotic cells could influence net cell death rate in early time-points post-impact and limit the timeframe when NAC can inhibit oxidative stress in lethally damaged cells. Nevertheless, inclusion of necrosis and apoptosis separately has been done in earlier computational models [28], and it could be included in the current modeling workflow when more experimental data become available.

Our simulations showed that immediately administered NAC treatment reduced acute cell death during the first hours after impact by inhibiting cellular oxidative damage (Fig 2A). With a high chondrocyte protection rate (*k*_c,dmg→c,h_ = 5.3. 10^−5^m^3^mol^−1^s^−1^), simulated cell viability increased from 46% to 77% in 2 days after impact compared to untreated cartilage in agreement with a 32 %-point increase seen in the *ex vivo* experiments [17] (Fig 4D). Similar efficiency of NAC was reported in human *ex vivo* experiments: 14 %-point increase in cell viability after 1-day NAC treatment (2 mM) [16] and increase of over 20 %-points after 7-day continuous (medium changed every 2-3 days) treatment of impacted cartilage compared to untreated samples [19]. Simulation results suggest that NAC could inhibit loss of cell viability throughout the tissue by counteracting cellular oxidative damage in superficial and deep zones of injured cartilage because of fast diffusion of NAC (small molecular size) to damaged regions.

After 1-, 4- and 12-hour post-impact treatment delays, our model predicted 69%, 55%, and 45% cell viability compared to 74±7%, 59±6%, and 39±6% cell viability, respectively, in the previous experiments [17] after a single administration of NAC (Fig. 4E). An earlier *ex vivo* human study reported that 7 days of continuous NAC treatment after a 24-hour delay could effectively increase cell viability above that of untreated samples after impact (0.59 J) [16]. This result could imply that with low impact energies, delayed administration of NAC after impact may still remain effective than currently suggested by our modeling framework (4 h). Although not simulated here, also estimating different impact energies and the associated different cell death rates affected by pro-inflammatory cytokines [28] is possible in the current modeling framework.

### 4.2. Proteoglycan content after impact

Without treatment, impact loading can trigger oxidative stress related proteoglycan degeneration in cartilage [16,20]. Our model showed 85% relative proteoglycan content at day 14 which was consistent with 88±10% relative proteoglycan content measured in the experiments [17] (Fig. 2C). The lowest proteoglycan content was observed in the superficial zone, which has been also reported in previous *ex vivo* studies of injuriously loaded cartilage [52] which could be driven by mechanical disruption of cartilage, loss of biosynthesis and increased proteolytic activity [16,52]. In our earlier model [28], we showed that over short periods, proteolytic enzyme production is more important to induce loss of proteoglycan content than cell death and impaired biosynthesis in injured and physiologically, cyclically loaded cartilage. This same mechanism is present in the current study which suggests decrease of proteoglycan biosynthesis was not the primary mechanism for the proteoglycan loss after impact. Hence, acute ROS inhibition by NAC could be important to effectively inhibit proteoglycan loss caused by catabolic cell reactions.

When NAC treatment was utilized immediately after impact (time 0) and after 4-hour delay, our simulations indicated that relative proteoglycan content in the impacted region (compared to the intact area) was increased by 10%-points and 3%-points compared to untreated cartilage at day 14, respectively (Fig. 2C). On average, the experiments reported an average increase of 22%-points in relative proteoglycan content at day 14 when treatment was not delayed [17]. Our modeling results suggest that after a 4-hour delay, treatment is no longer effective in reducing proteoglycan loss although it is still able to reduce cell death. Thus, our model suggests that acute inhibition of impact- and oxidative stress-related stimulation of catabolic enzymes (such as a disintegrin and metalloproteinase with thrombospondin-like motifs (ADAMTS)-4 and −5 [53–55]) may play an important role in preventing proteoglycan loss with NAC treatment [16,56]. However, in addition to reduced oxidative stress and cell damage, other mechanisms may also affect NAC-induced reduction in proteoglycan loss (Fig. 2C), such as an increase of proteoglycan biosynthesis by increased anabolic activities in cells [16,57], reduction of direct ROS-induced proteoglycan oxidation [58], and inhibited inflammatory response of chondrocytes [15,59]. Further research is needed to decipher the effect of each of these variables.

### 4.3 Limitations

Even though our computational modeling framework was able to replicate the experimentally observed cell viability and proteoglycan content, there are some limitations regarding its biomechanical and biochemical aspects. Our biomechanical simulations of the impact and the resulting strain distributions are dependent on the estimated material properties, cartilage thickness, and depth-dependent structural properties. Based on our preliminary tests, we observed that strain distribution during rapid impact loading was mostly influenced by collagen-related parameters (such as strain-dependent collagen fibril modulus and depth-dependent collagen orientation [32,33]), which were not analyzed by Martin et al. [17]. Also, the maximum shear strain threshold defining initiation of cell damage (40%) [22] and maximum cell damage (150%) [7] were based on earlier computational studies reporting cell death in areas exceeding the tissue-level strain-thresholds, but experiments have shown smaller cell-level strains causing cell death and damage [5]. These uncertainties may affect simulated strain distribution and initial cell damage distribution in cartilage, which may also influence the spatial NAC treatment effects predicted by our model. However, earlier studies have quantified cell deformation in near real-time during dynamic [60] and static [61] loading, and similar techniques combined with cell viability analysis may enable quantification of high strain-induced cell death after impact.

We included biochemical tissue degeneration and adaptation mechanisms that have the most experimental and computational support from literature. However, the resulting model remained simplified from *ex vitro*/*in vivo* conditions. The excluded mechanisms were inflammatory response of injured cartilage [6], release of damage-associated molecular patterns [62], degeneration caused by chondrocyte differentiation [63], altered NAC transport due to molecular charge of NAC [30] and decrease of NAC concentration via chemical reactions and uptake in the cells [64,65]. Since decrease of NAC concentration over time was not considered, our model may underestimate the efficiency of NAC because it would be smaller (degraded) concentration of NAC that actually causes the protection found against cell damage. In addition, our current model does not consider collagen degeneration, which can affect factors such as diffusion of different biomolecules [66]. Yet, our model offers a novel way to study temporal effects of degeneration and antioxidative treatments *ex vivo*, and augmenting the mechanisms in this baseline model would be the next step towards simulating cartilage degeneration and treatment-induced regeneration *in vivo*. Nevertheless, extensive experimental calibration and more quantitative data (for example gene expression, immunohistochemical analysis, and experiments with/without proteolytic enzyme-inhibitors to analyze NAC effects spatially and temporally) is needed to augment the current model with new mechanisms.

### 4.4 Future directions

In the future, this new modeling framework can be augmented with our previous modeling framework to simulate time-dependent cyclic loading and inflammation post-impact [28], time-dependent effects of treatment administration, sustained drug delivery, and multiple drug treatments to mitigate cartilage degeneration [67]. To advance this new modeling framework, the following biomechanical and biological data are needed: sample-specific material properties, fraction of damaged cells experiencing oxidative stress, location-specific reduction of damaged cells by NAC, fraction of live and dead cells, and quantitative data on proteoglycan content at several time-points post-impact. In addition, to analyze NAC uptake into cartilage and treatment effects in physiologically loaded cartilage, more data will be needed about possible lesions after impact, tissue structure and content, and activity of proteolytic enzymes within the cartilage. To gather these data, we will conduct new *ex vivo* experiments. Combining the calibrated cell-tissue level model into state-of-the-art joint-level degeneration models with patient-specific joint geometries, contact forces and inflammation [44,68], the model could then be used to guide the most optimal treatment strategies to mitigate PTOA progression.

## 5. Conclusions

We developed a novel computational modeling framework to study NAC treatment mechanisms on mitigating cartilage degeneration through overloading-driven cell damage. Our model successfully predicted reduced proteoglycan loss after NAC treatment following reduced impact-related cell damage and inhibited proteolytic activity. The developed modeling framework enhances understanding of the role of cell damage and oxidative stress on cartilage degeneration and time-dependent NAC treatment mechanisms post injurious loading of cartilage. Although definitive treatment has yet to be discovered for PTOA, our new modeling framework could aid the development of better treatment strategies. In the future, our modeling framework could help optimizing NAC dosage and timing for cartilage treatment to better inhibit cell death and cartilage degeneration after injurious loading.

## Acknowledgements

We acknowledge the support of University of Eastern Finland, Massachusetts Institute of Technology and University of Iowa to conduct this research.

## Supporting information captions

**S1 Structural, compositional, and material inputs for the modeling framework**.

**S2 Sensitivity analysis for peak impact force**.

**S3 Mesh convergence analysis**.

**S4 Proteoglycan degeneration in the biochemical model**.

**S5 Sensitivity analysis for aggrecanase release from damaged cells**.

## Notes

### Competing Interest Statement

The authors have declared no competing interest.

